# E3 ligase TRIM47 positively regulates endothelial activation and pulmonary inflammation through potentiating the K63-linked ubiquitination

**DOI:** 10.1101/2021.09.15.460553

**Authors:** Yisong Qian, Ziwei Wang, Hongru Lin, Tianhua Lei, Zhou Zhou, Weilu Huang, Xuehan Wu, Li Zuo, Jie Wu, Yu Liu, Ling-Fang Wang, Xiao-Hui Guan, Ke-Yu Deng, Mingui Fu, Hong-Bo Xin

## Abstract

Endothelial activation plays an essential role in the pathology of sepsis-induced acute lung injury, but the detailed regulatory mechanisms remain largely unknown. Here, we demonstrated that TRIM47, an ubiquitin E3 ligase of tripartite protein family, is highly expressed in vascular endothelial cells and is up-regulated during TNFα-induced endothelial activation. Knockdown of TRIM47 in endothelial cells prevents the transcription of multiple pro-inflammatory cytokines, reduces monocyte adhesion and the expression of adhesion molecules, and inhibits the secretion of IL-1β and IL-6 into the supernatant. By contrast, overexpression of TRIM47 promotes inflammatory response and monocyte adhesion upon TNFα stimulation. TRIM47 modulates the activation of NF-κB and MAPK signaling pathways during endothelial activation. Further experiment confirmed that TRIM47 interacts with TRAF2 and mediates K63-linked ubiquitination. In addition, TRIM47-deficient mice are more resistant to lipopolysaccharide-induced acute lung injury and death, due to attenuated pulmonary inflammation. Taken together, our studies suggest that TRIM47 promotes pulmonary inflammation and injury at least partly through potentiating the K63-linked ubiquitination of TRAF2, which in turn activates NF-κB and MAPK signaling pathways to trigger inflammatory response in endothelial cells.

## Introduction

Acute Lung injury (ALI) comprise a uniform response of the lung to inflammatory or chemical insults and is therefore commonly generated by systemic illness including sepsis or trauma, infection with pathogens and toxic gas inhalation ^1^. Despite a great deal of effort has been devoted to target the immune response to infection and adjunct approaches, there are few effective therapeutic strategies due to the emergence of new pathogens such as the global pandemic of novel coronavirus pneumonia, as well as the continued rise of drug resistance ^2^.

The pulmonary endothelium is critically implicated in the pathogenesis of ALI as a main target of circulating cells and humoral mediators under injury ^1^. The interaction between endothelial cells and leukocytes is a key step in the development of ALI. Leukocyte adhesion to endothelial cells and migration across endothelial cells are mediated by the interaction of complementary adhesion molecules on leukocytes and endothelial cells. The increased expression or release of endothelial cell adhesion molecules is a hallmark of endothelial cell activation ^3, 4^. Upon the access of leukocytes into the lung parenchyma, they can release inflammatory mediators to destroy pathogens, but the over-activated immune response have potential to result in the imbalance of the pro-inflammatory and anti-inflammatory mechanisms, triggering “cytokine storm” and subsequent tissue damage ^2, 5^. Considering the critical role of endothelial response during ALI, strategies that targeting endothelial components, including cell surface receptors ^6^, signaling pathways, transcriptional networks, and endothelial cell gene products, have been recently proposed to attenuate endothelial activation and improve endothelial dysfunction ^7^.

Tripartite motif-containing (TRIM) proteins, a subfamily of E3 ubiquitin ligases, have been widely involved in many physiological processes including cell proliferation and differentiation, innate immunity and autophage ^8^. Multiple TRIM proteins have been found to participate in innate immunity through positive or negative regulation of cytokines, toll like receptors, pattern recognition receptors, intracellular signaling pathways and transcription factors ^9^. Nemours studies have focused on the roles and regulatory mechanisms of TRIM in immune cells ^10-14^, there are relatively few studies on TRIM proteins involved in the regulation of endothelial inflammation. It has been shown that TRIM28 is abundant in endothelial cells, and interfering with TRIM28 expression has anti-inflammatory and anti-angiogenic phenotypes ^15^. However, the precise roles of TRIM family members in regulating of endothelial function remain largely unknown.

TRIM47 was originally found in brain astrocytomas and was named GOA (gene overexpressed in astrocytoma). It is prominently located in nucleus and its LXXLL motifs are thought to be closely related to nuclear receptor binding ^16^. Emerging evidence showed that TRIM47 have functions on tumorigenesis and progression ^17-19^, viral resistance processes ^20^, and cerebral ischemia-reperfusion injury ^21^. A recent genome-wide association study showed that the SNPs of TRIM47 and TRIM65, located in the adjacent position of human chromosome 17, are closely related to white matter hyperintensities, which are thought to reflect ischemic damage to the small deep cerebral vessels ^22^. These findings suggested that TRIM47 and TRIM65 may coordinately or independently participate in the regulation of cerebrovascular injury. Our previous work demonstrated that TRIM65, as an E3 ubiquitin ligase, selectively targets VCAM-1 and promote its ubiquitination and degradation, thus reducing lung inflammation and damage caused by sepsis ^23^, but the role of TRIM47 in endothelial inflammation remains to be elucidated.

In the present study, we investigate the effects of TRIM47 in endothelial activation in an *in vitro* model of inflammation induced by TNFα. The global TRIM47 knockout mice was constructed to confirm its role in ALI. The E3 ubiquitin ligase activity and substrate molecules of TRIM47, as well as the signaling pathways involved were also explored.

## Materials and methods

### Reagents

Human recombinant TNFα and LPS (from *Escherichia coli* O111:B4) were purchased from Sigma. VCAM-1 (sc-13160), ICAM-1 (sc-1511-R), β-actin (sc-47778), Histone H2A (sc-10807) antibodies were from Santa Cruz Biotechnology. TRIM47 antibody (26885-1-AP) was purchased from Proteintech. Phospho-p65 (3033), p65 (8242), IκBα (4812), phospho-IκBα (2859), phospho-IKKα/β (2078), IKKα (11930), IKKβ (8943), phospho-JNK (4668), JNK (9252), phospho-ERK1/2 (4370), ERK1/2 (4695), phospho-p38 (4511), p38 (8690), Flag (8146 and 2368) and α-tubulin (2125) antibodies were purchased from Cell Signaling Technology. Ubiquitin (ab7780), ubiquitin (K48, ab140601), and ubiquitin (K63, ab179434) antibodies were purchased from Abcam.

### Cell culture, infection and treatment

Human Umbilical Vein Endothelial Cells (HUVEC) and THP-1 cells were purchased from Lonza Walkersville Inc. HUVECs were cultured in EGM medium according to the manufacturer instruction, and used for experiment in less than eight passages. THP-1 was cultured in RPMI-1640 medium supplemented with 10% FBS and 2-mercaptoethanol to a final concentration of 0.05 mM. RAW264.7, EA.hy926, bEnd.3, and HEK293 cells were purchased from ATCC and cultured in DMEM supplemented with 10% FBS. U251, HeLa, MDA-MB-23, 3T3-L1 and A549 cell lines were from National Collection of Authenticated Cell Cultures (Shanghai, China) and cultured in DMEM supplemented with 10% FBS. The hCMEC/D3 cell line was purchased from BeNa Culture Collection (Beijing, China) and cultured in RPMI-1640 medium supplemented with 10% FBS. HL-1 cardiac muscle cell line was obtained from Sigma Aldrich and cultured in Claycomb medium supplemented with 100 μM norepinephrine, 4 mM l-glutamine and 10% FBS. The siRNA target sequence was selected in Human TRIM47 gene (GenBank accession NM_033452.2) as follows: siRNA1: 5’-TGAAGCTCCCAGGGACTATTT-3’, and siRNA2: 5’-TACTGGGAGGTGGAGATTATC-3’. TRIM47 siRNA was constructed into the lentivirus expression vector pLV[shRNA]-EGFP:T2A:Puro-U6. A universal sequence was used as a negative control for RNA interference. Human TRIM47 gene was constructed into pLV[Exp]-EGFP:T2A:Puro-EF1A vector to obtain the expression lentiviral vector. The viral particles were produced by third generation packaging in 293T cells and Lentiviral stocks were concentrated using ultracentrifugation. HUVECs (5 × 10^4^/ml) were prepared and infected at a Multiplicity of Infection (MOI) of 50 with negative control, TRIM47 siRNA1, TRIM47 siRNA2 or TRIM47 overexpression lentiviruses for 16 h at 37°C in the presence of 10 mg/ml polybrene. The cultures were then washed and cultured in fresh medium for 72 h. GFP expression was detected to calculate the infection efficiency. Then, cells were treated with 10 ng/ml TNFα for indicated times, and mRNAs or proteins from those cells were extracted and detected.

### RNA isolation and QPCR

Total tissue or cellular RNA was isolated using TRIzol reagents, according to the manufacturer’s instructions (Life Technologies, CA, USA). One microgramme of total RNA was reverse-transcribed using a One Step PrimeScript™ RT-PCR Kit (Takara, Liaoning, China) with a thermocycler. The mRNA levels were determined by SYBR Green dye using an ABI 7500 sequence detection system with a reaction mixture that consisted of SYBR Green 2×PCR Master Mix (Applied Biosystems, CA, USA), cDNA template, and the forward and reverse primers. Primer sequences are listed in Table S1. The PCR protocol consisted of 40 cycles of denaturation at 95 °C for 15 s followed by 60 °C for 1 min to allow extension and amplification of the target sequence. Data were analyzed using ABI 7500 sequence detection system software. The amount of mRNA was normalized to GAPDH using the 2^−ΔΔCT^ method. The results were from three independent experiments performed in triplicate.

### Protein isolation and western blot

Tissue extracts and whole-cell lysates were prepared in radioimmunoprecipitation assay buffer (Thermo Scientific) supplemented with 1 mM PMSF. Nuclear and cytoplasmic protein fractions from cells were extracted by Nuclear-Cytosol Extraction Kit (Applygen Technologies Inc, Beijing, China), according to the manufacturer’s instructions. Fifty micrograms protein per sample was loaded in each lane and separated by sodium dodecyl sulfate-polyacrylamide gel electrophoresis (SDS-PAGE) and transferred to nitrocellulose membranes (Pall Corporation, USA) in Tris-glycine buffer (48 mM Tris, 39 mM glycine, pH 9.2) containing 20% methanol. The membranes were blocked with skimmed milk for 1 h, washed in Tris buffered saline containing 0.1% Tween-20 (TBST), and incubated with primary antibodies overnight at 4 °C. After washing in TBST for three times, nitrocellulose membranes were incubated for 1 h at room temperature with the horseradish peroxidase conjugated IgG (1:5000; Santa Cruz Biotechnology, Inc, CA, USA). The bands were visualized by the SuperSignal West Pico Chemiluminescent Substrate Trial Kit (Pierce, Rockford, IL, USA). The immunodetected protein bands were then analyzed using ChemiDoc XRS system with Quantity One software (Bio-Rad, Richmond, CA, USA).

### Immunocytochemistry

At the end of the treatment, cells were rinsed with phosphate-buffered saline (PBS) three times, fixed with 4% paraformaldehyde for 30 min at room temperature and permeabilised in 0.1% Triton X-100 for 10 min. An incubation in 5% bovine serum albumin (BSA) in PBS for 1 h was performed to prevent antibody non-specific binding. The cultures were incubated with primary antibodies overnight at 4 °C. After incubation with primary antibodies, cells were incubated with fluorescein isothiocyanate (FITC)-conjugated goat anti-rabbit IgG (Alexa 488; 1:1000; invitrogen) and the nuclei were stained with DAPI. Immunostained cells were examined under a fluorescence microscope (Olympus IX71, Tokyo, Japan).

### Ubiquitination assay with Co-IP

HUVECs were transfected with Flag-tagged TRIM47 plasmid or empty vector using electroporation. The protease inhibitor MG132 (10 mM; Sigma-Aldrich) was added 4 h before harvest. At 24 h posttransfection, cells were lysed in CelLytic M Cell lysis buffer with protease inhibitors, phosphatase inhibitors, NEM, and ubiquitin aldehyde. For endogenous IP, HUVECs were harvested after 15 min-TNFα (10 ng/mL) exposure. The immune complexes were collected by incubation (2 h, 4°C) with protein G-agarose (Sigma). Co-IP assays were performed by using Pierce™ Protein G Agarose (Thermo Fisher) followed the manufacturer’s instruction. After extensive washing, the electrophoresis loading buffer was added to the complexes and incubated for 5 minutes at 95°C. Immunoprecipitated proteins were resolved by SDS-PAGE and analyzed by immunoblotting with indicated antibodies.

### Generation of TRIM47 knockout mice

To define the physiological role of TRIM47 in vivo, we have obtained the mice with heterozygous TRIM47-targeted allele by using CRISPR/Cas9 to remove all exons of TRIM47 gene (Fig. 5A). The mice with homozygous TRIM47-targeted alleles were generated by interbreeding. The mice were created in C57BL/6 background. Genotyping was done with the following primers: Trim47-F: 5’-GGTAAACACAGTCGCTAAGAGGTCAAA-3’, Trim47-R: 5’-TGGTCTAGGGATGCCAGGGTTCT-3’ and Trim47-Wt/He-F: 5’-AGTCAGAGTGAGCAGGCAGGAGAATA-3’ (Fig. 5B). Wild type and TRIM47 knockout mice were housed in the Animal Centre of Institute of Translational Medicine, Nanchang University, with a 12 h light-dark cycle, optimal temperature and humidity, filtered water, and appropriate nutrient feed. All procedures related to the care of animals were performed according to the National Institutes of Health Guide for the Care and Use of Laboratory Animals. All experimental protocols were approved by Institutional Animal Care and use Committee of Nanchang University. The lung tissue extracts were examined by western blot analysis and the results showed that TRIM47 protein was abolished in the homozygous targeted allele (Fig. 5C), which generated the global knockout mouse (designated as TRIM47^−/−^).

### LPS challenge in mice

Age-matched mice (7-9 weeks) were randomly assigned to control or experimental groups. Wild type and TRIM47^−/−^ mice underwent an intraperitoneal injection of LPS (15 mg/kg) to induce lethal endotoxic shock. The control group received injections of the equivalent volume of 0.9% NaCl solution. After injection, the mice were closely monitored for general condition and survival for 7 days.

### Measurements of inflammatory cytokines

The concentrations of TNFα, IL-1β and IL-6 in the serum, and IL-1β and IL-6 levels in the supernatant of HUVECs were measured using the specific ELISA kits according to the manufacturer’s instructions (Neobioscience Technology Co., Ltd., Shenzhen, China). Absorbance at 450 nm wave length was measured, and the protein concentration was determined by interpolation on to absorbance curves generated by recombinant protein standards using iMark™ Microplate Absorbance Reader (Bio-Rad).

### Histological analysis

The right upper lungs were removed after 24 h of LPS challenge and were fixed in 4% phosphate-buffered paraformaldehyde. The 4-μm paraffin tissue sections were cut and stained with H&E as previously described ^24^. Photomicrographs were taken by a light microscope (Olympus BX51). Lung injury was evaluated by an independent pathologist who was blinded to the grouping, taking into account haemorrhage in the lung tissue, alveolar congestion, edema, infiltration of macrophages and neutrophils, and morphological changes in the alveolar wall.

### Measurement of lung wet/dry weight ratio

To evaluate the magnitude of pulmonary edema, the wet-to-dry weight ratios at 24 h after LPS challenge were determined. The left lung tissue samples were weighed immediately after removal (wet weight) and then subjected to desiccation in an oven at 50 °C until a stable dry weight was achieved after 72 h. The ratio of the wet/dry weight was then calculated.

### Statistics

Statistical analysis was performed with GraphPad Prism software (GraphPad, San Diego, CA). Data were expressed as mean ± SD. For comparison between two groups, the unpaired Student’s t-test was used. For multiple comparison, one way ANOVA followed by Turkey’s post hoc analysis was used. A value of p < 0.05 was considered significant.

## Results

### TRIM47 is highly expressed in vascular endothelial cells

The mRNA and protein levels of TIRM47 were detected in various tissues of mice. Results showed that TRIM47 mRNA was highly expressed in heart, lung, kidney and epididymal white adipose tissue (eWAT, Fig. 1A). TRIM47 protein levels were abound in heart, lung, stomach and testis (Fig. 1B). The immunohistochemistry demonstrated that TRIM47 had a high expression in lung, kidney tubules, heart and eWAT, moderate expression in brain, stomach, skin and colon, and low expression in liver, testis, spleen and thymus (Fig. S1). In particular, an obvious positive staining was observed in the vascular lining of multiple tissues, as indicated by the arrows, including lung, brain, subcutaneous tissue and colon (Fig. 1C). Next, we detected TRIM47 expression in different cells. Real-time PCR and western blot results revealed that TRIM47 was specifically expressed in human umbilical vein endothelial cells (HUVEC and EA.hy926) and brain microvascular endothelial cells (bEnd.3), whereas TRIM47 showed low expression in monocyte/macrophages (Fig. 1D and 1E). In addition, we measured the mRNA levels of 56 TRIM family members in HUVECs and hCMECs, respectively. TRIM47 exhibited a moderate expression in both cells (Fig. 1F and 1G). These results suggested that TRIM47 is highly expressed in vascular endothelial cells.

**Fig. 1.**
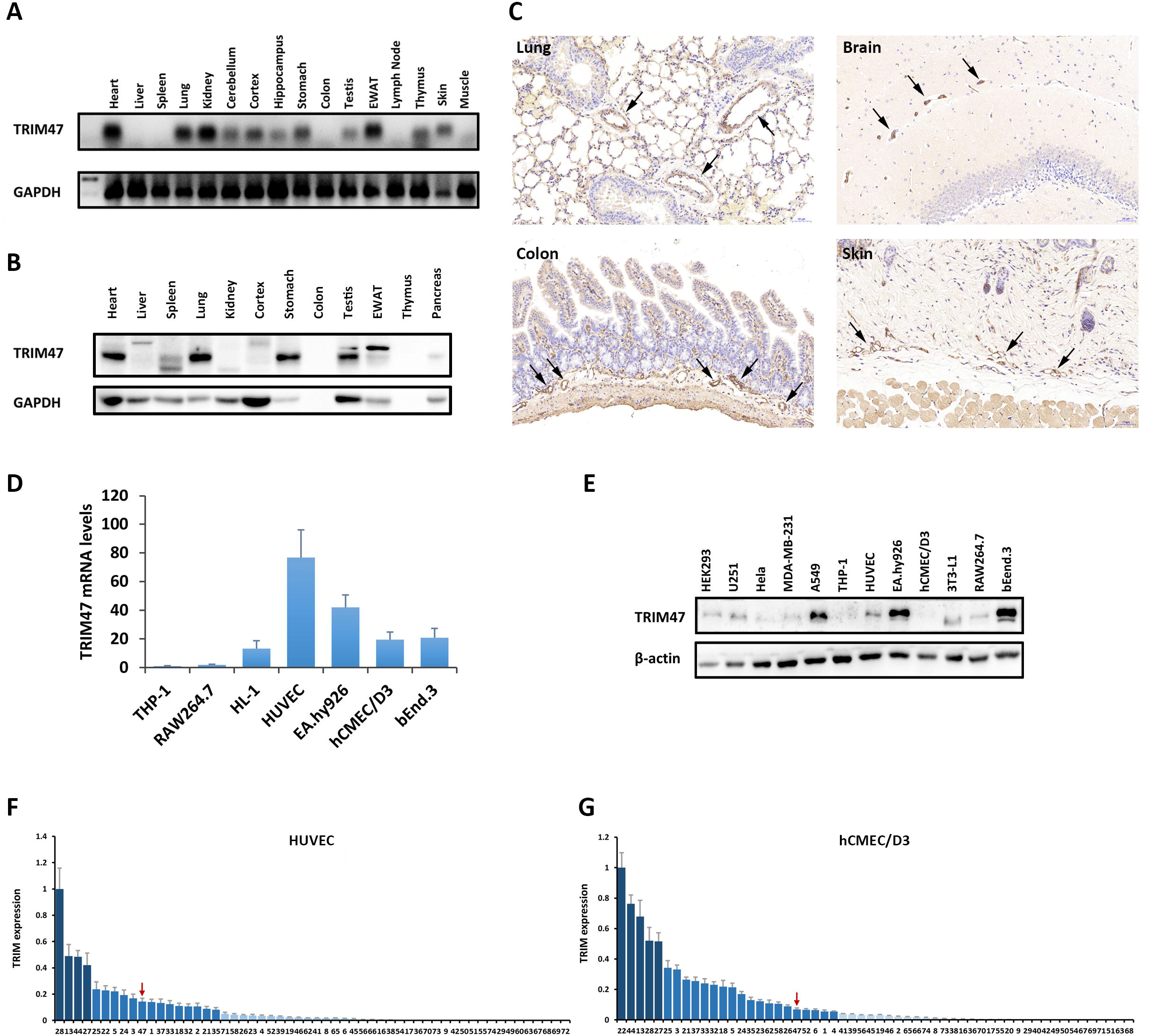
TRIM47 is highly expressed in vascular endothelial cells. (A) The mRNA levels of TIRM47 in various tissues of mice were detected by real-time PCR and visualized by agarose gel electrophoresis. (B) The protein levels of TIRM47 were detected in various tissues of mice by western blot. (C) The immunohistochemistry showed positive staining of TRIM47 in the vascular lining of multiple tissues. (D) TRIM47 mRNA expression in different cells detected by real-time PCR. (E) The protein levels of TRIM47 in different cells detected by western blot. (F) The mRNA levels of 56 TRIM family members in HUVECs and hCMEC/D3 detected by real-time PCR.

### TRIM47 is induced by inflammatory stimulation in endothelial cells

We further investigated the expression changes of TRIM47 following exogenous stimuli. Result showed that the mRNA and protein levels of TRIM47 were significantly up-regulated by LPS, H_2_O_2_ and TNFα (Fig. 2A-C). The expression of TRIM47 was increased since 2 h after TNFα stimulation and peaked at 12 h (Fig. 2C). Similarly, TRIM47 expression was also up-regulated after TNFα challenge in bEnd.3 cells upon various stimulation, including inflammation and hypoxia (Fig. S2A-D). However, TRIM47 was not sensitive to TNFα challenge but remarkable decreased by LPS exposure in macrophages (Fig. S2E-H). The localization of TRIM47 was measured by western blot in cytosolic and nuclear fractions, respectively. The nucleus contains a prominent proportion of TRIM47, which was significantly up-regulated after TNFα exposure. The cytosolic TRIM47 was also induced after TNFα stimulation (Fig. 2D). The immunocytochemistry further confirmed that TRIM47 mainly but not solely located in the nucleus of HUVECs. One-hour stimulation of TNFα resulted in an increase in fluorescence density, which was partly resumed at 8 h of TNFα exposure (Fig. 2E). TRIM47 also responded to LPS challenge in HUVECs (Fig. S3A), and was up-regulated in hCMEC/D3 following TNFα and LPS incubation (Fig. S3B and S3C). The expression profile of TRIM47 indicated that it may play a role during inflammatory stimulation.

**Fig. 2.**
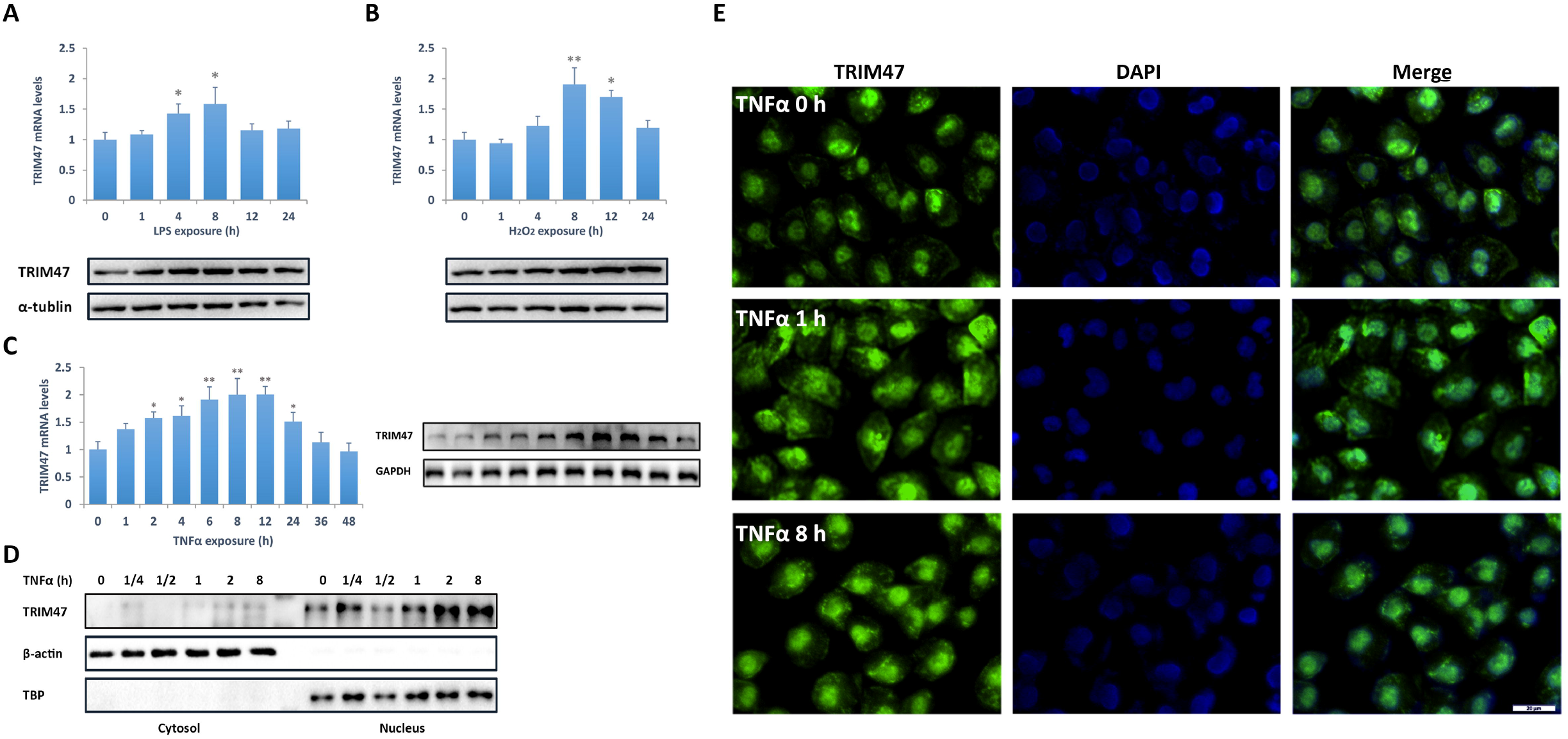
TRIM47 is induced by inflammatory stimulation in endothelial cells. TRIM47 mRNA and protein levels in HUVECs after (A) LPS, (B) H_2_O_2_ and (C) TNFα stimulation were examined by real-time PCR and western blot, respectively. (D) The translocation of TRIM47 following TNFα treatment was measured by western blot in cytosolic and nuclear fractions. (E) The distribution of TRIM47 in HUVECs was detected by immunocytochemistry in the absence or presence of TNFα.

### TRIM47 promotes TNFα-induced endothelial activation

The siRNA and overexpression vectors were introduced to explore the role of TRIM47 in endothelial activation. Knockdown of TIRM47 significantly reduced TNFα-induced mRNA expression of multiple adhesion molecules and pro-inflammatory cytokines (Fig. 3A), which were elevated in TRIM47-overexpressed HUVECs. TRIM47 siRNAs obviously decreased the protein levels of ICAM-1 and VCAM-1, and suppressed THP-1 adhesion to HUVECs, whereas overexpression of TRIM47 had opposite effects (Fig. 3C and 3D). TRIM47 siRNAs also suppressed the secretion of pro-inflammatory cytokines IL-1β and IL-6 into the supernatant (Fig. 3E and 3F). By contrast, overexpression of TRIM47 promotes the production of pro-inflammatory cytokines (Fig. 3G and 3H). In addition, knockdown of TRIM47 remarkably inhibited cell proliferation and migration induced by TNFα in HUVECs (Fig. S4). These results suggested that TRIM47 is a positive regulator of endothelial activation.

**Fig. 3.**
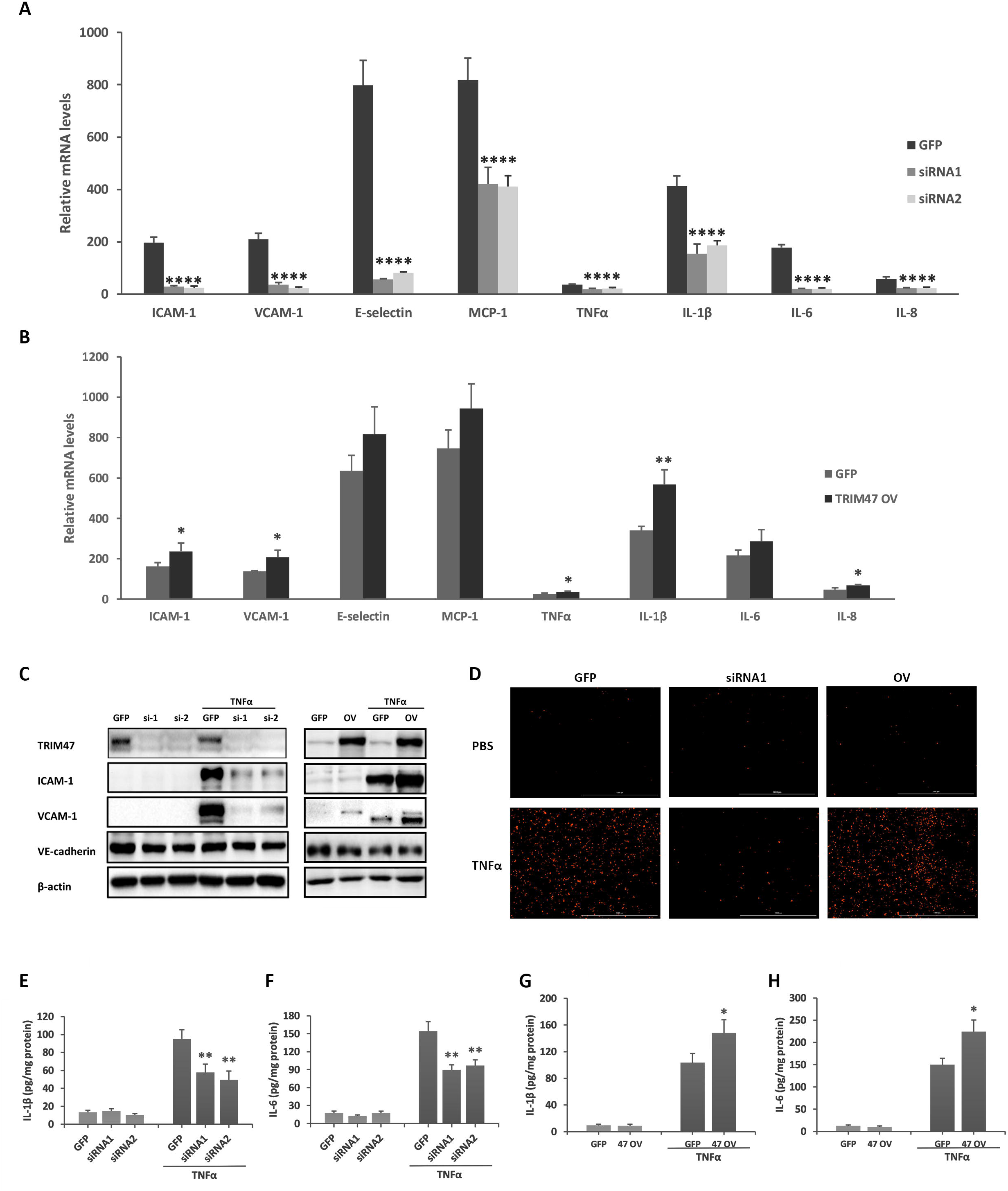
TRIM47 promotes inflammatory response in endothelial cells. The mRNA expression of multiple adhesion molecules and pro-inflammatory cytokines was detected by real-time PCR in (A) TRIM47 siRNA- and (B) overexpression vector-treated HUVECs. (C)The protein levels of ICAM-1 and VCAM-1 were measured by western blot in TRIM47 siRNA- and overexpression vector-treated HUVECs. (D) TRIM47 siRNA or expression vector was transfected into HUVECs, and the transfected cells were incubated with TNF-α or PBS for 8 h and then co-cultured with Zombiered-labeled THP-1 cells for 1 h. The adhesion of THP-1 was observed under a fluorescence microscope. The contents of (E) IL-1β and (F) IL-6 in siRNA-treated cells, and (G) IL-1β and (H) IL-6 in overexpression vector-treated cells were measured by ELISA.

### TRIM47 modulates NF-κB and MAPK pro-inflammatory signaling pathways

The potential signaling pathway involved in TRIM47-mediatied endothelia activation was investigated, including NF-κB and MAPK. Results showed that Knockdown of TRIM47 significantly inhibited the phosphorylation of IκBα, IKKα/β and p65 subunit, and prevented the degradation of IκBα (Fig. 4A), whereas overexpression of TRIM47 promotes the activation of NF-κB signaling pathway (Fig. 4B). TRIM47 siRNAs also suppressed the activation of JNK and p38 MAPK signal pathways but had no obvious effect on ERK (Fig. 4C). Overexpression of TRIM47 further activated JNK and p38 signal pathways after TNFα stimulation (Fig. 4D). These results suggested that TRIM47 mediates endothelial activation at least partly through NF-κB and MAPK activation.

**Fig. 4.**
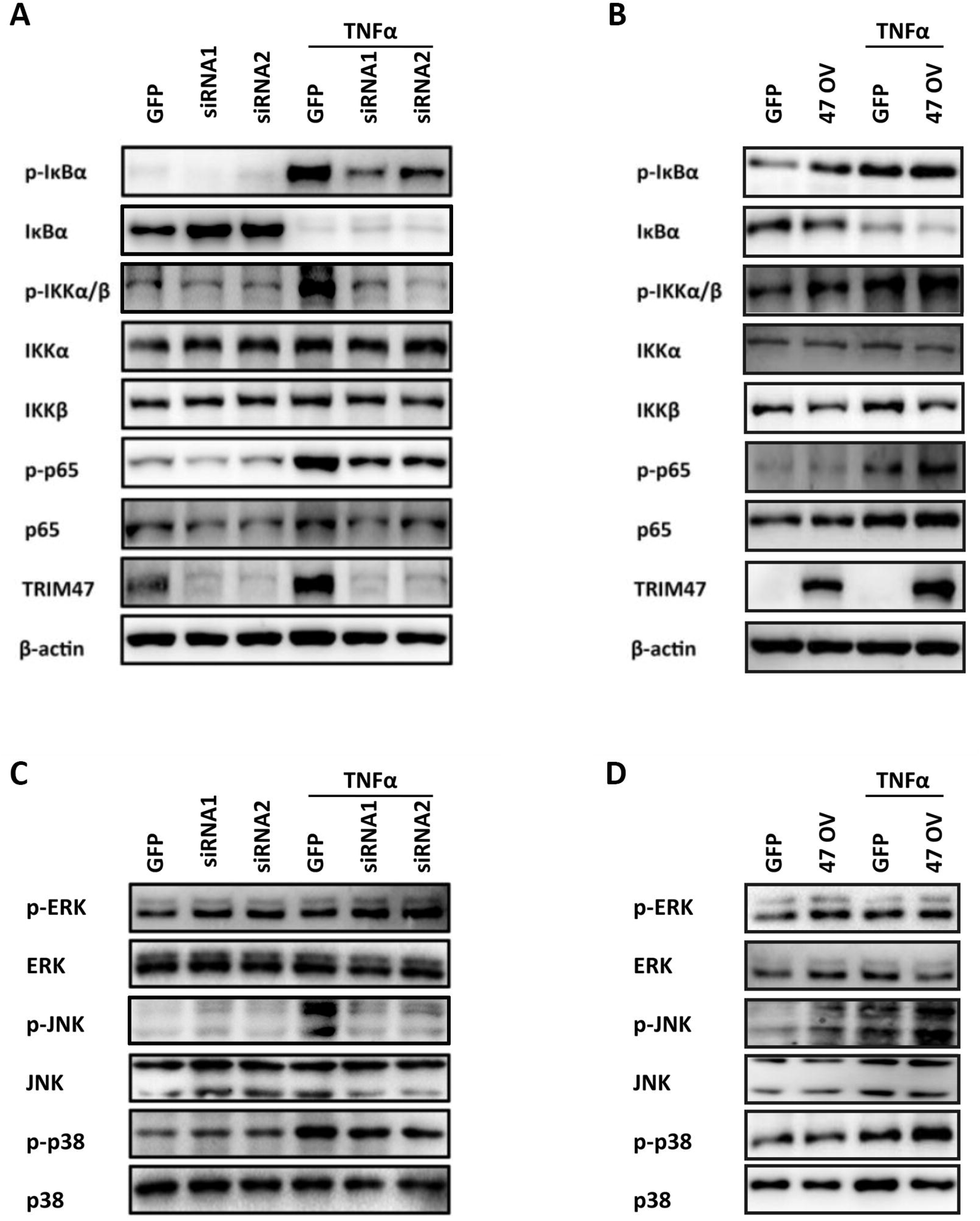
TRIM47 modulates endothelial activation through NF-κB and MAPK signaling pathway. The activation of NF-κB was detected by western blot in (A) TRIM47 siRNA- and (B) overexpression vector-treated HUVECs. The activation of MAPK, including ERK, JNK and p38 signaling pathways was detected by western blot in (C) TRIM47 siRNA- and (D) overexpression vector-treated HUVECs.

### TRIM47 mediates K63-linked Ubiquitylation and interacts with TRAF2

Emerging evidence showed that that TRIM proteins mediated K48- or K63-linked ubiquitination to activate NF-κB signaling pathway in response to exogenous stimulation ^12, 25^. Therefore, the ubiquitination pattern of TRIM47 involved in endothelial activation was analyzed. We observed that TNF-α induced total ubiquitination and K63-linked ubiquitination of TRIM47 in HUVECs by Co-immunoprecipitation with the TRIM47 antibody (Fig. 5A). HUVECs were transfected with Flag or Flag-TRIM47 vector for 48 h. Co-immunoprecipitation was performed with the Flag antibody. The exogenous assay further confirmed that TRIM47 was involved in K63-linked ubiquitination rather than K48 (Fig. 5B). Tumor necrosis factor receptor-associated factor 2 (TRAF2) is a key adaptor molecule in TNFR signaling complexes that promotes downstream signaling cascades, such as NF-κB and MAPK activation ^26^, whereas TRAF6 is the major transducer of IL-1 receptor/TLR signaling ^27^. It has been reported that the K63-linked polyubiquitin chains could be attached to TRAF2, serving as a scaffold to recruit TAK1, TAB1, and TAB2. The active TAK1 further phosphorylates the MAPKs and IKK complex to initiate MAPK and NF-κB cascades ^28^. We therefore examined the possible binding proteins of TRIM47 involved in this signal pathway. As shown in Fig. 5C, TRIM47 bind with TRAF2 but not TRAF6. TRIM47 did not interact with the downstream proteins including TAK1, IKKγ and IκBα. In addition, TRIM47 did not induce the degradation of IκBα, a classical target of K48-linked ubiquitination (Fig. S5). These results suggest that TRIM47 regulated NF-κB and MAPK activation by enhancing the K63-linked ubiquitination of TRAF2.

**Fig. 5.**
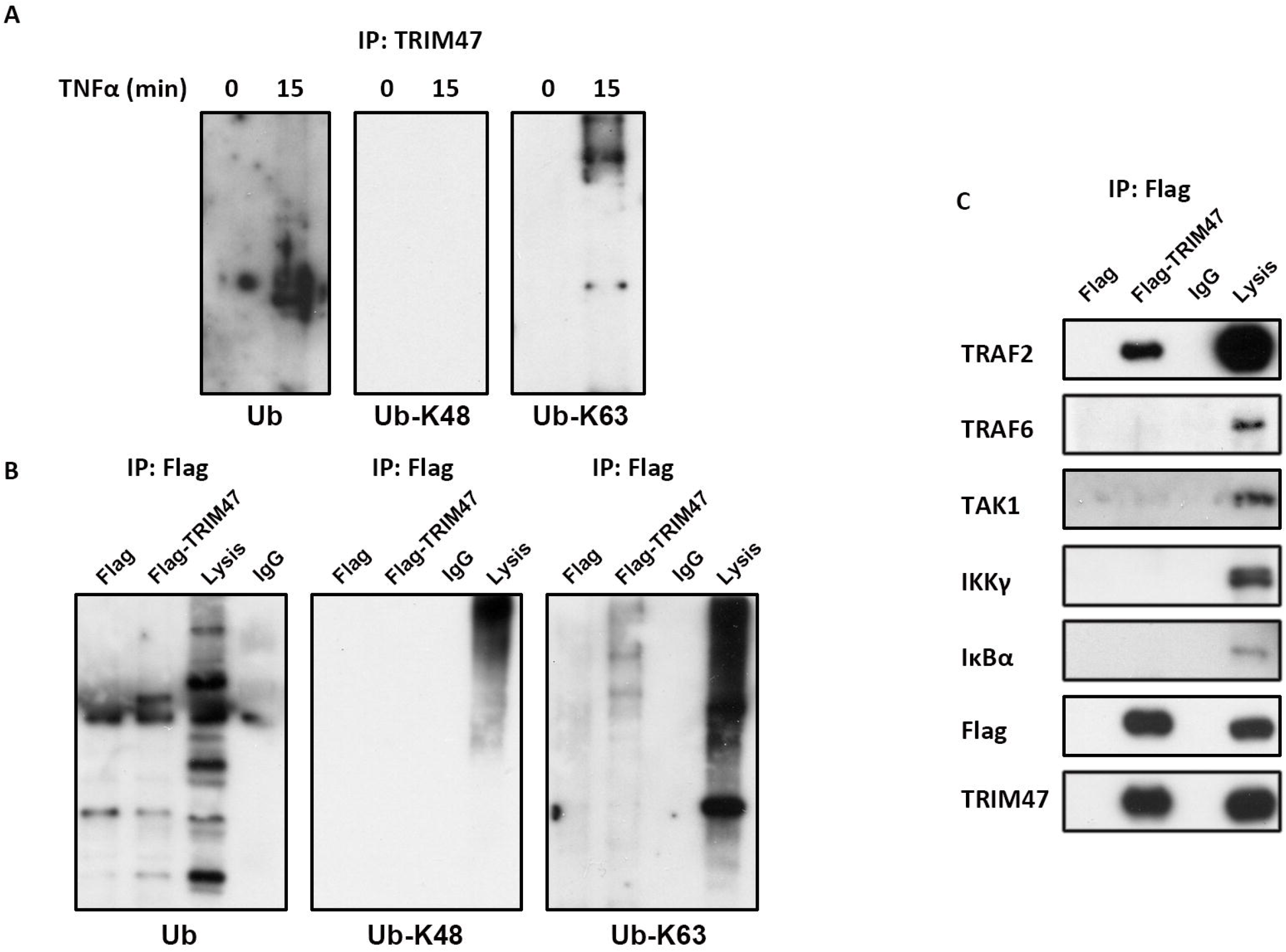
TRIM47 mediates K63-linked Ubiquitylation and interacts with TRAF2. (A) HUVECs were treated with 10 ng/ml TNF-α for 0 and 15 min. Cell lysates then were immunoprecipitated using TRIM47 antibody, followed by western blot analysis using the indicated antibodies. (B) HUVECs were transiently transfected with empty vector or TRIM47 plasmid for 48 h. Whole-cell lysates were immunoprecipitated with Flag antibody and the precipitates were immunoblotted with Ub, TRAF2, TRAF5, TAK1, IKKγ, and IκBα antibodies.

### TRIM47 deficiency alleviates acute lung injury and inflammatory response in LPS-challenged mice

We produced the global TRIM47 knockout mice to investigate its role in systemic inflammatory response and organ injury (Fig 6A-C). The TRIM47 KO mice had no significant changes in viscera index (Table. S2) or histology (Fig. S6) compared with the WT mice. After 24 h of LPS injection, the TRIM47 KO mice showed reduced pulmonary edema compared with the WT animals (Fig. 6D). In addition, TRIM47 deficiency improved the survival rate of mice after LPS challenge (Fig. 6E). HE staining showed that there were no significant differences in lung histology between WT and TRIM47 KO mice under normal conditions. A significant tissue damage appeared in the lungs of WT mice, including neutrophil infiltration, alveolar wall thickening, hemorrhage, alveolar edema and alveolar disruption. The histological changes were significantly alleviated in TRIM47 KO mice (Fig. 6F).

**Fig. 6.**
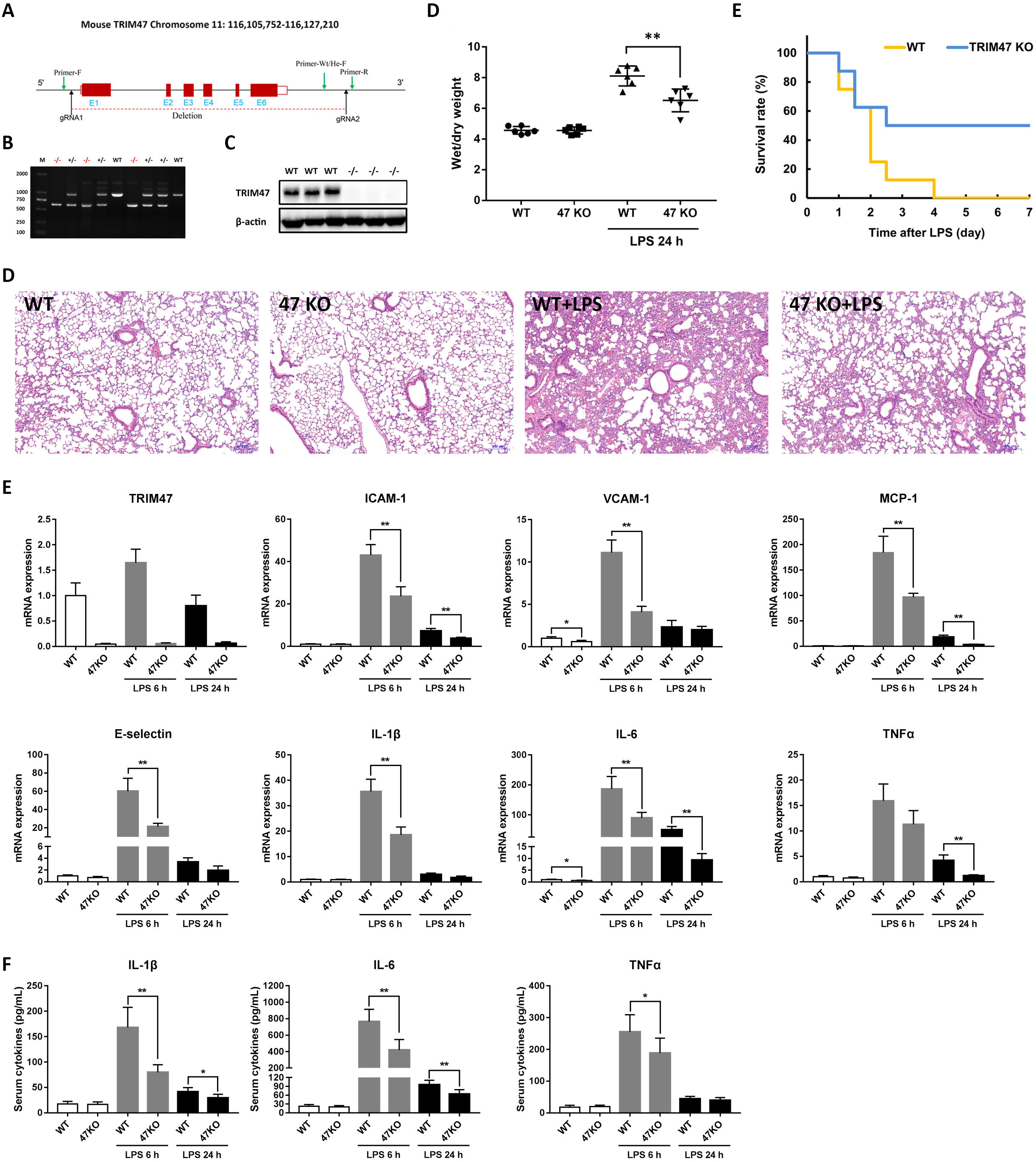
TRIM47 deficiency reduces endotoxemia-induced acute lung injury and pulmonary inflammation in mice. (A) Schematic strategy of generation of TRIM47 knockout mice. (B) Genotyping of TRIM47^+/+^, TRIM47^+/™^, and TRIM47^™/™^ mice. (C) Western blot analysis for TRIM47 in lungs from TRIM47^+/+^ and TRIM47^™/™^ mice. (D) Pulmonary edema was represented as lung wet-to-dry ratio. (E) Survival rate of mice challenged with 15 mg/kg LPS (i.p.). (F) The histological changes in lungs after LPS challenge were examined by HE staining. (G) The mRNA levels of TRIM47 and various pro-inflammatory cytokines in the lung were measured by real-time PCR. (F) The content of IL-1β, IL-6 and TNFα in the serum was assayed by ELISA.

TRIM47 and various pro-inflammatory cytokines in the lung and serum were measured after 6 h and 24 h following LPS injection, respectively. The mRNA levels of pro-inflammatory cytokines were much lower in TRIM47 KO mice than that in WT mice (Fig. 6G). The knockout mice also had lower levels of TNFα, IL-1β and IL-6 content in the serum (Fig. 6H). These results indicated that TRIM47 deficiency attenuates systemic inflammatory and acute lung injury during LPS challenge.

## Discussion

Here we demonstrated that TRIM47 play crucial roles in TNFα-induced endothelial activation. As a potential E3 ubiquitin ligase, TRIM47 interacts with TRAF2 and mediates K63-linked ubiquitination, activating NF-κB and MAPK signaling pathways. In LPS challenged mice, TRIM47 deficiency significantly reduced pulmonary inflammation and tissue damage. These results strongly suggested that TRIM47 is a positive regulator of endothelial activation.

TRIM47 was originally found to be overexpressed in astrocytoma tumor cells and astrocytes of fetal brain, with prominent nuclear staining, but virtual absence in mature astrocytes. The mRNA levels of TRIM47 in normal tissues were low except for in kidney ^16^. Here we showed that TRIM47 has relatively high expression in heart, lung, kidney and eWAT by detecting the mRNA and protein levels respectively. The immunohistochemistry also demonstrated its extensive distribution in these tissues. Interestingly, the vascular endothelial cells from different tissues exhibited remarkable positive staining of TRIM47. The samples from cell cultures further confirmed its prominent expression in various endothelial cells. Therefore TRIM47 may be identified as a specifically expressed protein in endothelial cells where may take its functions.

A various TRIM proteins have been proved to be implicated in innate immunity and inflammatory response. Our previous work showed that TRIM47 is down-regulated upon TNFα exposure in THP1-derived macrophages ^29^. According to the present study, TRIM47 was significantly induced by multiple stimuli, including TNFα, LPS, hypoxia and oxidative stress in endothelial cells, indicating that TRIM47 may be a sensor in response to exogenous stimulation. In agreement with the previous report ^16^, we found that TRIM47 was prominent located in the nucleus of HUVECs and was expressed in cytoplasm upon TNFα challenge. These result suggested TRIM47-mediated endothelial activation may depend on its cytoplasm location, although its role in nucleus remain unclear.

Different from the anti-inflammatory effect of TRIM65, another constitutive expressed protein in endothelial cells ^23^, TRIM47 promotes inflammatory response both *in vitro* and *in vivo*. The *in vivo* results further confirmed that the pro-inflammatory phenotype primarily depend on endothelial activation rather than macrophages. Since the previous GWAS results suggested that both TRIM47 and TRIM65 contribute to the cerebral vessels injury ^22^, we may suppose that TRIM47 and TRIM65 are critically involved in the regulation of endothelial inflammation and injury by different mechanisms.

Proteins covalently modified with K48-linked poly-ubiquitin are targeted for proteasomal degradation, whereas proteins covalently modified with K63-linked poly-ubiquitin generally become functionally activated ^9^. Recent work showed that TRIM proteins positively or negatively mediate NF-κB activation primary through K48- or K63-linked poly-ubiquitin. For example, TRIM5 interacts with TAK1-containing kinase complex and positively regulate NF-κB activation by mediating K63 poly-ubiquitin chain synthesis ^12^. TRIM25 promote NF-κB activation by enhancing the K63-linked ubiquitination of TRAF2 and bridging the interaction of TRAF2 and TAK1 or IKKβ ^28^. TRIM14 enhances NF-κB activation in endothelial cells via directly binding to NEMO and promotes the phosphorylation of IκBα and p65, which is dependent on its K63-linked ubiquitination ^30^. TRIM47 is structurally similar to TRIM25, with a Ring-finger domain, a B2 box and its associated coil-coiled region at the N terminus, and a PRY domain at the C terminus ^8^. Functionally like TRIM25, TRIM47 also interacts with TRAF2 and promotes MAPK and NF-κB activation through K63-linked ubiquitination. These findings provide the underlying mechanisms by which TRIM47 specifically meditates endothelial activation (Fig. 7).

**Fig. 7.**
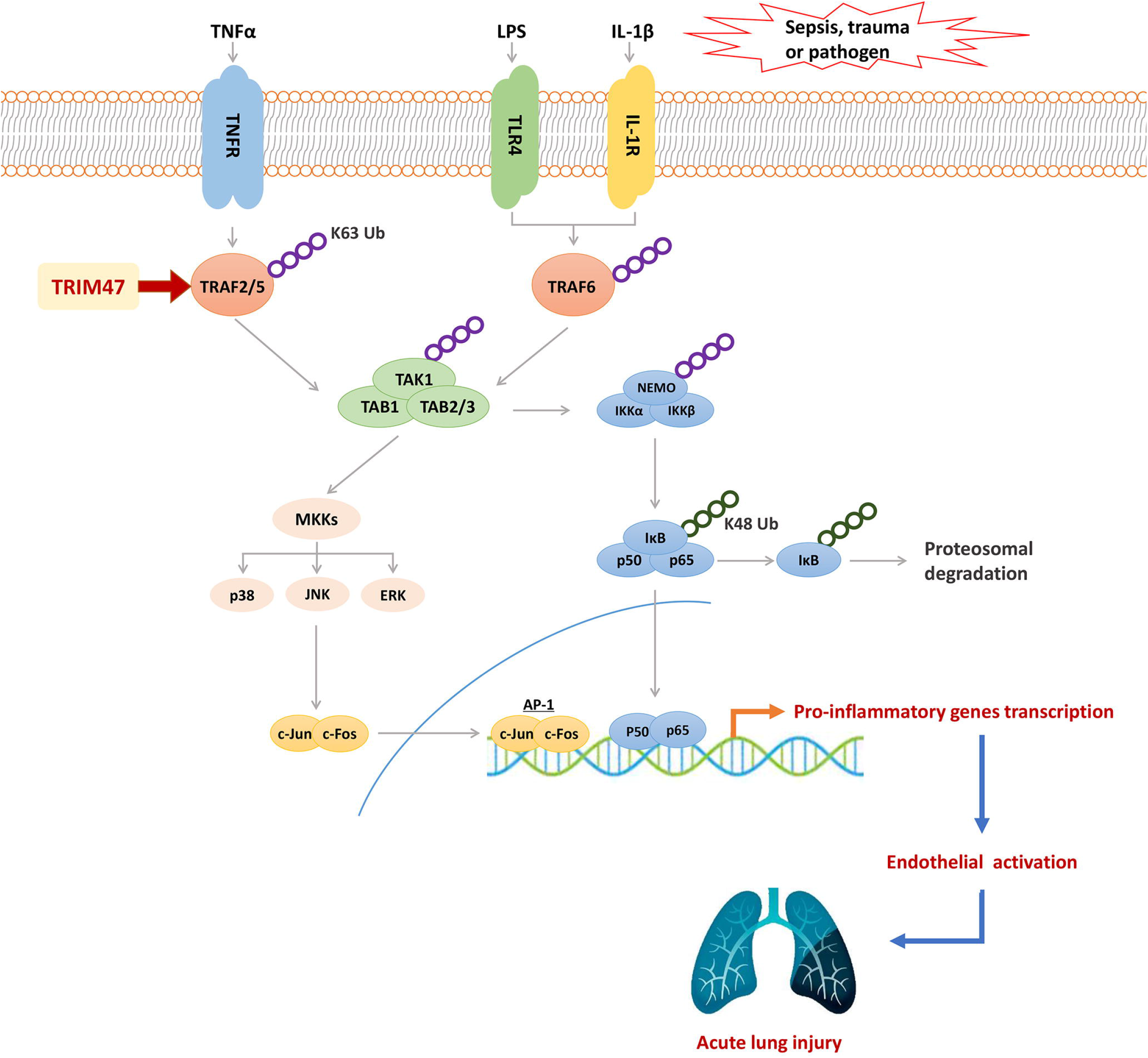
The schematic diagram of the mechanism by which TRIM47 promotes TNFα-induced endothelial inflammation and acute lung injury.

To sum up, we identified a novel endothelial activation factor TRIM47, which mediates inflammatory response in endothelial cells and promote inflammation and tissue damage during ALI through endothelial TRAF2-MAPK/NF-κB pro-inflammatory axis. Currently there remain no compounds targeting TRIM proteins at the laboratory or clinical level but it is important to develop inhibitors of TRIM proteins for their use as a therapeutic tools in multiple diseases ^8^. TRIM47 would be an attractive drug target for endothelial inflammation and ALI. Further detailed analysis of TRIM47 is needed for its use for effective therapy and to eliminate side effects.

## Acknowledgement

This work was supported by the National Natural Science Foundation of China (82070080 and 81860020 to YQ, 81873659 to HBX, and 81760140 and 81970256 to KYD), Foundation for the National Institutes of Health (1R15AI138116 to M.F), and the financial support provided by China Scholarship Council (201906825031 to YQ).

## CRediT authorship contribution statement

Yisong Qian, Ziwei Wang, Hongru Lin, Tianhua Lei, Zhou Zhou, Weilu Huang, Xuehan Wu, Li Zuo, Jie Wu, Yu Liu: Investigation, Methodology. Ling-Fang Wang, Xiao-Hui Guan and Ke-Yu Deng: Resources. Yisong Qian: Writing-original draft, Conceptualization, Supervision. Mingui Fu and Hong-Bo Xin: Conceptualization, Supervision, Writing-review & editing.

## Declaration of competing interest

The authors declare that they have no known competing financial interests or personal relationships that could have appeared to influence the work reported in this paper.

## Notes

### Competing Interest Statement

The authors have declared no competing interest.

