## supplemental material for "E3 ligase TRIM47 positively regulates endothelial activation and pulmonary inflammation through potentiating the K63-linked ubiquitination"

**Tab. S1 Primers used in real-time PCR reactions**

| Gene | Forward primer (5'→3') | Reverse primer (5'→3') |
|---|---|---|
| **HumTRIM47** | GAGGGTGCTGTGTCCTATCAACT | CGATAATCTCCACCTCCCAGTAGT |
| **HumVCAM-1** | AGGAAGGCAGTTCTGTGAATATGAC | CCTGCTCCACAGGATTTTCG |
| **HumICAM-1** | CGTTGCCTAAAAAGGAGTTGCT | GCTATCTTCTTGCACATTGCTCAGT |
| **HumE-selectin** | CTGGACTCTCCCTCCTGACATT | ACAAATTTCTTTGCTTTCCGTAAGC |
| **HumMCP-1** | CGCCTCCAGCATGAAAGTCT | GGAATGAAGGTGGCTGCTATG |
| **HumTNFα** | TCTTCTCGAACCCCGAGTGA | GGCCCGGCGGTTCA |
| **HumIL-1β** | CCACAGACCTTCCAGGAGAATG | ATCCCATGTGTCGAAGAAGATAGG |
| **HumIL-6** | TCCAGGAGCCCAGCTATGAA | GAGCAGCCCCAGGGAGAA |
| **HumIL-8** | GCCAAGGAGTGCTAAAGAACTTAGA | TGGTCCACTCTCAATCACTCTCA |
| **HumGAPDH** | CAGGGCTGCTTTTAACTCTGGT | GATTTTGGAGGGATCTCGCT |
| **MusTRIM47** | GGCCCCCAGGGATTACTTC | CCAAAAAGCTGCAGGAACTTG |
| **MusVCAM-1** | GACTCCATGGCCCTCACTTG | GCGTTTAGTGGGCTGTCTATCTG |
| **MusICAM-1** | CACCCCGCAGGTCCAAT | CAGAGCGGCAGAGCAAAAG |
| **MusE-selectin** | GGGACCCAACTGTGAGCAA | ACGGGTGGGAGCAGTTCAG |
| **MusMCP-1** | CTCTCTCTTCCTCCACCACCAT | AGCCGGCAACTGTGAACAG |
| **MusTNFα** | ATCCGCGACGTGGAACTG | ACCGCCTGGAGTTCTGGAA |
| **MusIL-1β** | CTACAGGCTCCGAGATGAACAAC | TCCATTGAGGTGGAGAGCTTTC |
| **MusIL-6** | GTGCAATGGCAATTCTGATTGT | GGTAGCATCCATCATTTCTTTGTATCT |
| **MusGAPDH** | ACATGGCCTCCAAGGAGTAAGAA | GGGATAGGGCCTCTCTTGCT |

**Table. S2 Viscera index in WT and TRIM47 knockout mice (n = 6).**

| | **WT** | **TRIM47 KO** |
|---|---|---|
| Heart | 0.585±0.021 | 0.566±0.010 |
| Liver | 6.814±0.383 | 7.324±0.372 |
| Spleen | 0.330±0.060 | 0.277±0.016** |
| Lung | 0.805±0.020 | 0.859±0.027 |
| Kidney | 0.837±0.032 | 0.822±0.030 |
| Brain | 1.320±0.020 | 1.388±0.052 |

** P <0.01 vs WT.

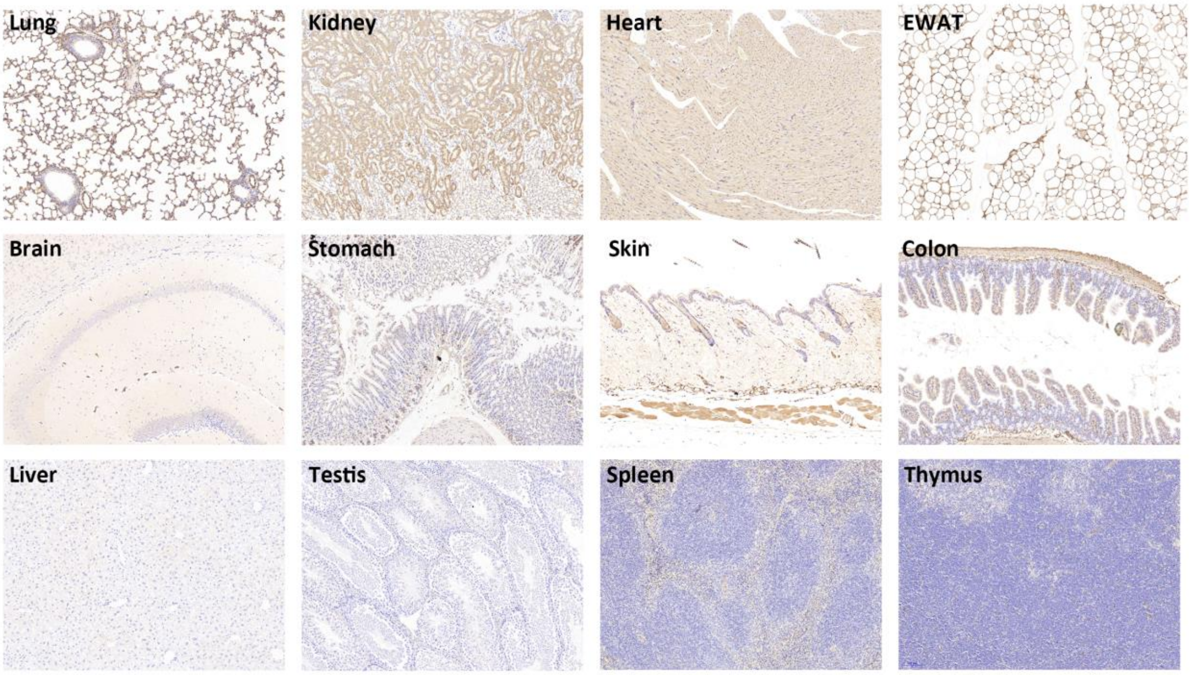


**Fig. S1 TRIM47 expression in different tissues detected by immunohistochemistry.**

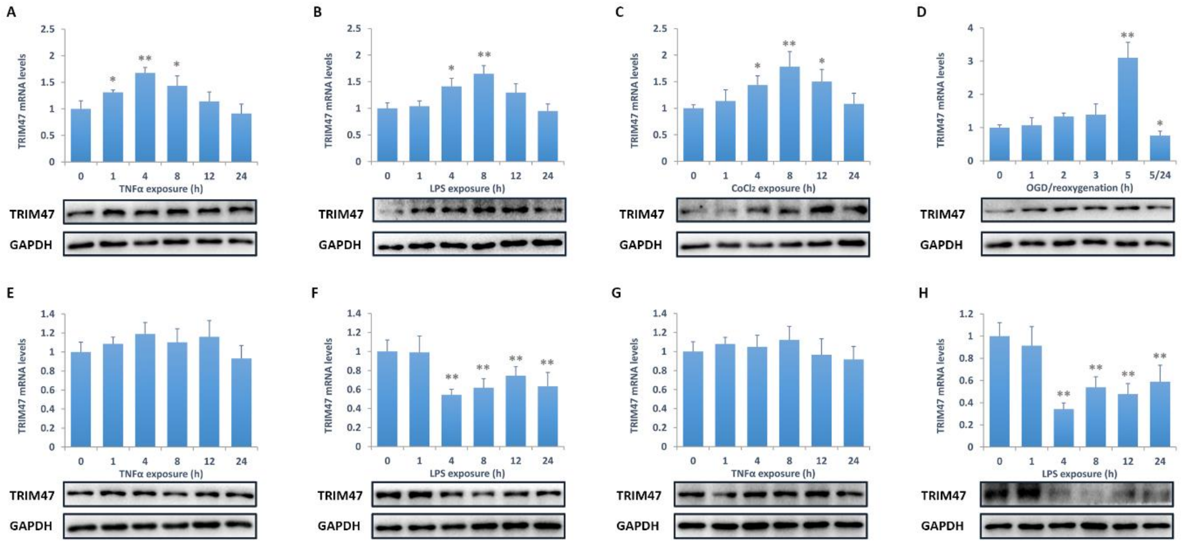


**Fig. S2 TRIM47 expression after exogenous stimuli.** TRIM47 mRNA and protein levels were determined by real-time PCR and western blot respectively. The mouse brain microvascular endothelial cells bEnd.3 were challenged with (A) TNFα, (B) LPS, and hypoxia induced by (C) $CoCl_2$ and (D) oxygen glucose deprivation/reoxygenation (OGD/R). The peritoneal macrophages were exposed to (E) TNFα and (F) LPS, and RAW264.7 macrophages were exposed to (G) TNFα and (H) LPS. *P <0.05, **P < 0.01 vs the control.

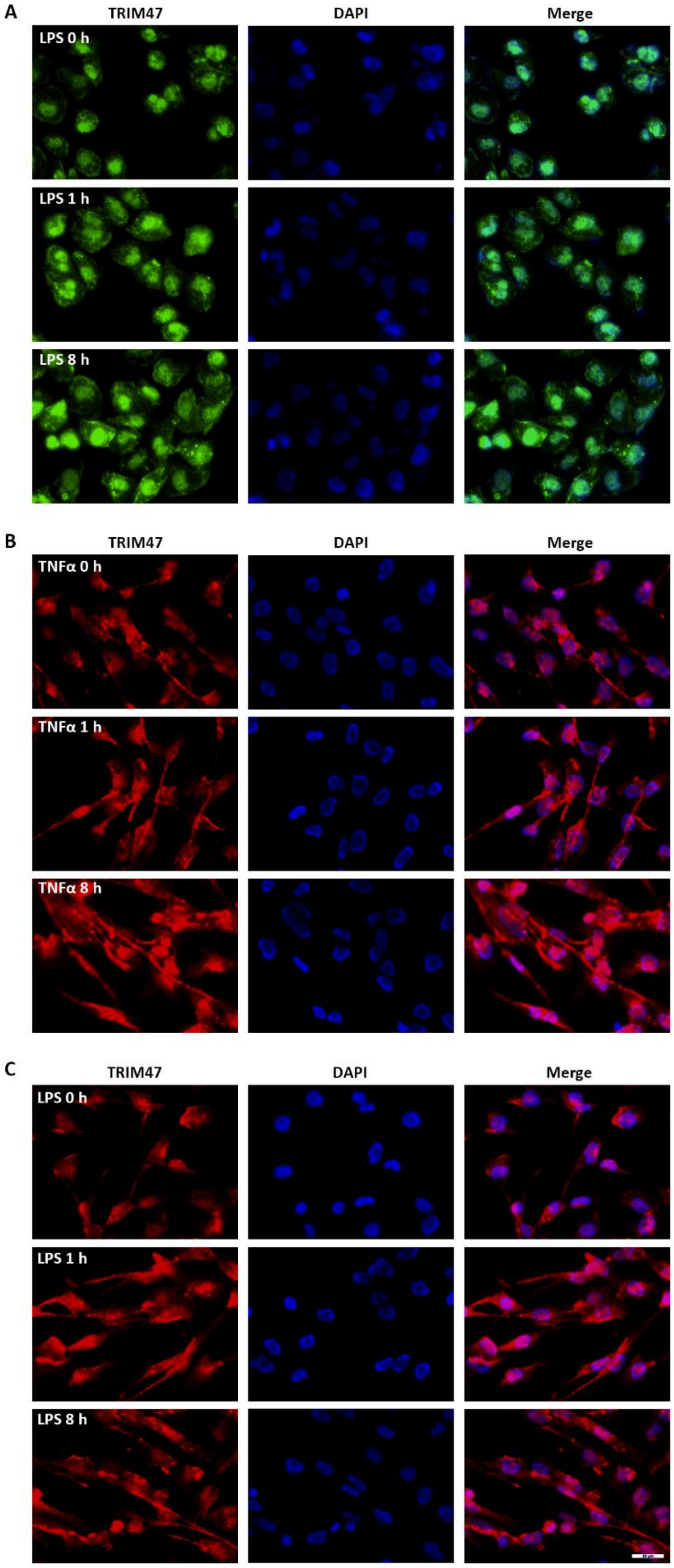


**Fig. S3 TRIM47 distribution after inflammatory stimulation.** Cell cultures were immunostained at 1 h and 8 h after inflammatory stimulation. (A) HUVECs were treated with LPS (100 ng/mL). HCMEC/D3 cultures were treated with (B) TNFα and (C) LPS.

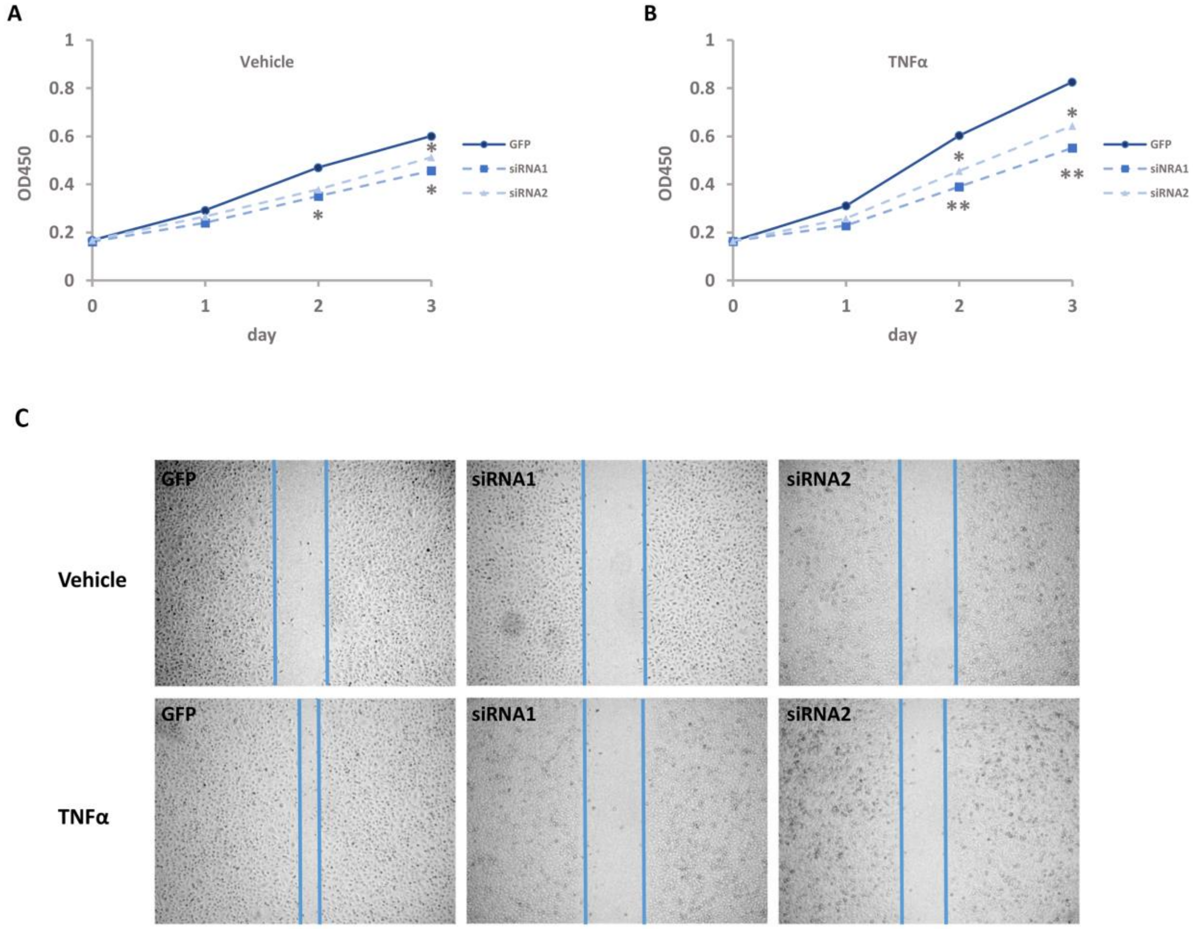


**Fig. S4 Knockdown of TRIM47 inhibits cell proliferation and migration induced by TNFα.** The proliferation of HUVECs were determined by CCK8 method (A) in the absence and (B) in the absence of TNFα. (C) Cell migration was observed by scratch.

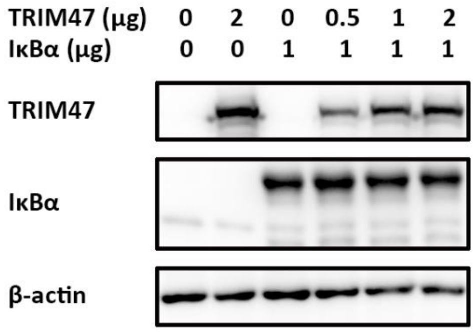


**Fig. S5 TRIM47 dose not induce IκBα degradation.** The TRIM47 and IκBα vectors were co-transfected into HEK293 cells for 24 h. The protein levels of TRIM47 and IκBα were determined by western blot.

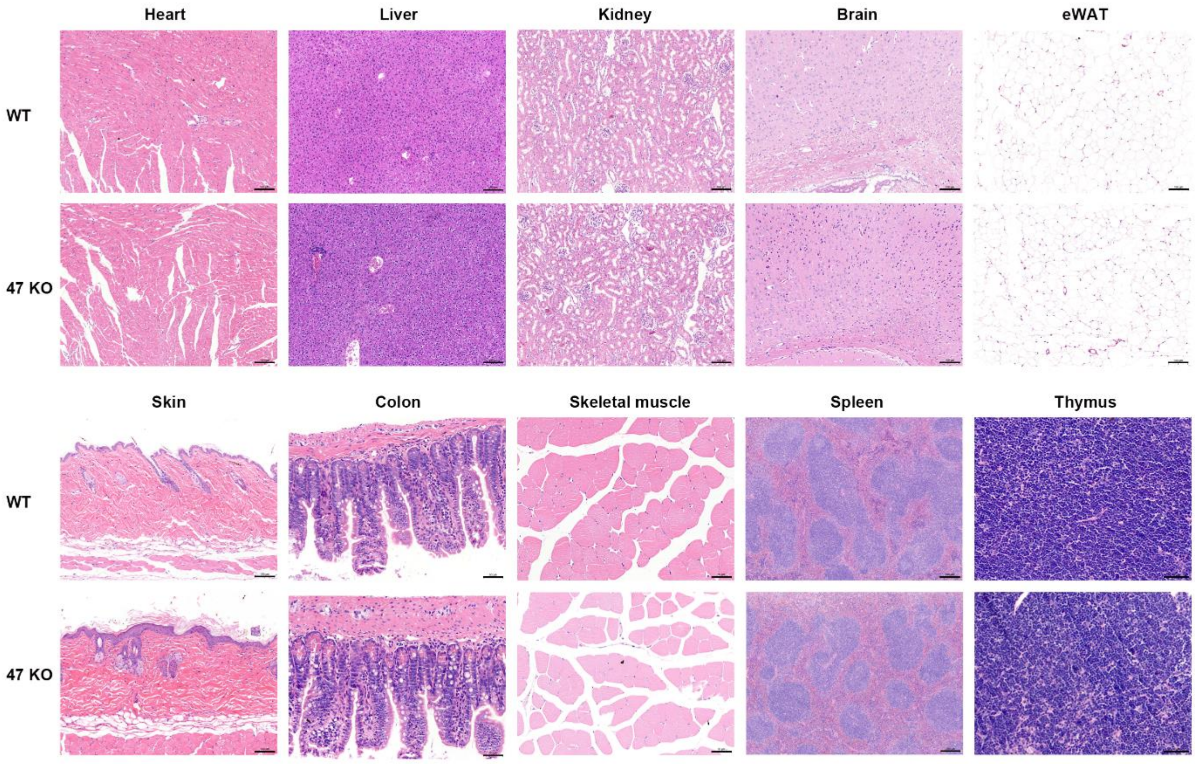


**Fig. S6 Histology measurement by HE staining in WT and TIRM47 KO mice.**
